# Rhythmic chromatin interactions with lamin B1 reflect stochasticity in variable lamina-associated domains during the circadian cycle

**DOI:** 10.1101/584011

**Authors:** Annaël Brunet, Frida Forsberg, Philippe Collas

**Author notes:** These authors contributed equally to this work (Annaël Brunet, Frida Forsberg).

## Abstract

Many mammalian genes exhibit circadian expression patterns concordant with periodic binding of transcription factors, chromatin modifications and chromosomal interactions. Here, we report periodic interactions of chromatin with nuclear lamins, suggesting rhythmic associations with the nuclear lamina. Entrainment of the circadian clock is accompanied in mouse liver by a gain of lamin B1-chromatin interactions, followed by oscillations in these interactions at hundreds of lamina-associated domains (LADs). A subset of these oscillations exhibit distinct 12, 18, 24 or 30-h periodicity in our dataset, and affect one or both LAD borders or entire stand-alone LADs. However, most LADs are conserved during the circadian cycle, and periodic LADs are seldom occurrences rather than dominant features of variable LADs. Periodic LADs display oscillation asynchrony between 5’ and 3’ LAD borders, and are uncoupled from periodic gene expression within or in vicinity of these LADs. Accordingly, periodic genes, including central clock-control genes, are often located megabases away from LADs, suggesting residence in a transcriptionally permissive environment throughout the circadian cycle. Autonomous oscillatory associations of the genome with nuclear lamins provide new evidence for rhythmic spatial chromatin configurations. Nevertheless, our data suggest that periodic LADs reflect stochasticity in lamin-chromatin interactions underlying chromatin dynamics in the liver during the circadian cycle. They also argue that periodic gene expression is by and large not regulated by rhythmic chromatin associations with the nuclear lamina.

## Introduction

Thousands of mammalian genes exhibit autonomous oscillatory patterns of expression concordant with the circadian (24 h) rhythm (Hastings et al. 2018). The circadian rhythm is governed by central and peripheral clocks, respectively in the nervous system and in individual organs, controlled by transcriptional and translational negative feedback loops (Takahashi 2017). The core clock is regulated by the CLOCK and BMAL1 transcription factors (TFs) which drive expression of clock-controlled genes including *Per, Cry, Nr1d1/Nr1d2* (encoding REV-ERB alpha/beta proteins, respectively) and *Ror* genes (encoding ROR alpha/beta/gamma), by binding to E-boxes in their promoters. The PER-CRY repressor complex inhibits activity of CLOCK-BMAL1, lowering transcription of *Per* and *Cry* and generating a negative feedback loop. RORs and REV-ERBs respectively act as activators and repressors of *Arntl* (also called *Bmal1*) and other clock genes, driving their rhythmic transcription. Stability of PER and CRY proteins is regulated by post-translational modifications leading to their time-dependent degradation, enabling a new cycle of CLOCK-BMAL1-driven gene expression (Takahashi 2017).

Circadian binding of TFs and chromatin modifiers to promoters and enhancers generates rhythmic chromatin modifications and remodeling (Katada and Sassone-Corsi 2010; Feng et al. 2011; Koike et al. 2012; Masri et al. 2014; Zhang et al. 2015; Kim et al. 2018). In mouse liver, histone H3K4me3 oscillates at promoters of circadian genes (Vollmers et al. 2012; Aguilar-Arnal et al. 2015), while rhythmic H3K4me1 and H3K27ac levels define oscillating enhancers (Koike et al. 2012; Vollmers et al. 2012; Fang et al. 2014; Takahashi 2017). Recruitment to chromatin of the sirtuin SIRT1, a histone deacetylase (HDAC) involved in circadian control of metabolism (Nakahata et al. 2008; Masri et al. 2014), is under influence of oscillatory levels of metabolites (Aguilar-Arnal et al. 2015) and provides a molecular link between metabolism, chromatin and circadian rhythms. Periodic recruitment of HDAC3 to chromatin also regulates circadian rhythms (Feng et al. 2011). These oscillatory cistromes and chromatin modifications raise the possibility that other chromatin-linked processes also show rhythmic patterns. Indeed, periodic short- and long-range promoter-enhancer interactions regulate and connect circadian liver gene expression networks (Aguilar-Arnal et al. 2013; Xu et al. 2016; Kim et al. 2018; Mermet et al. 2018). Thus, circadian-dependent changes in chromatin topology contribute to shaping the nuclear landscape (Yeung and Naef 2018).

Dynamic interactions of chromatin with the nuclear lamina, a meshwork of A-type lamins (lamins A/C; LMNA, LMNC) and B-type lamins (LMNB1, LMNB2) at the nuclear periphery (Burke and Stewart 2013) also constitute one mechanism of regulation of gene expression (van Steensel and Belmont 2017). Lamina-associated domains (LADs) are typically heterochromatic and most LADs are conserved between cell types (Peric-Hupkes et al. 2010). However, other LADs are more variable and are altered during differentiation (Peric-Hupkes et al. 2010; Rønningen et al. 2015; Robson et al. 2016; Poleshko et al. 2017) or by mutations in nuclear lamins (Paulsen et al. 2017). It remains unclear however to what extent variable (v)LADs arise and disappear as a consequence of regulatory mechanisms or through random interactions of chromatin with nuclear lamins. Whether individual loci or broader domains such as LADs display oscillatory interactions with the nuclear lamina has also to our knowledge not been addressed.

Scarce evidence links the nuclear envelope to circadian gene expression. HDAC3, a component of the clock negative feedback loop (Shi et al. 2016) and a regulator of lamina-associated genes (Demmerle et al. 2013), interacts with the inner nuclear membrane proteins TMPO (also called lamina-associated polypeptide 2 beta) (Somech et al. 2005) and emerin (Demmerle et al. 2013). The clock regulators SIRT1 and SIRT6 deacetylases interact with LMNA (Liu et al. 2012; Ghosh et al. 2015) at the nuclear lamina, where they modulate histone acetylation and gene expression. *In vitro*, BMAL1 expression seems to be modulated by MAN1, another protein of the inner nuclear membrane, through MAN1 binding to the *BMAL1* promoter (Lin et al. 2014). Lastly, PARP1 and CTCF have been shown in a human colon cancer cell line to mediate cyclic interactions of specific genes with the nuclear lamina, promoting oscillating transcription (Zhao et al. 2015). These observations suggest contributions of the nuclear envelope to the regulation of circadian gene expression. However, whether chromatin exhibits genome-wide scale rhythmic associations with the lamina has not been examined.

Here, we determined whether chromatin exhibits periodic associations with nuclear lamins after entrainment of the circadian clock. We show that periodic lamin B1-chromatin interactions, although detectable, are not a dominant feature of variable LADs during the circadian cycle in mouse liver, and are uncoupled from periodic gene expression.

## Results

### Hepatic genes exhibit distinct periodic transcript levels in the liver

To entrain the circadian clock, mice were subjected to 24 h fasting and refed *ad libitum* from circadian time CT0. Livers were collected every 6 h until CT30, as well as from non-synchronized mice 18 h before the fasting period (Fig. 1A). Entrainment of the clock by fasting/refeeding was confirmed by the cyclic expression of the core clock genes *Clock, Arntl* (*Bmal1*), *Cry1, Per1* and *Nr1d1* (Fig. 1B).

**Figure 1.**
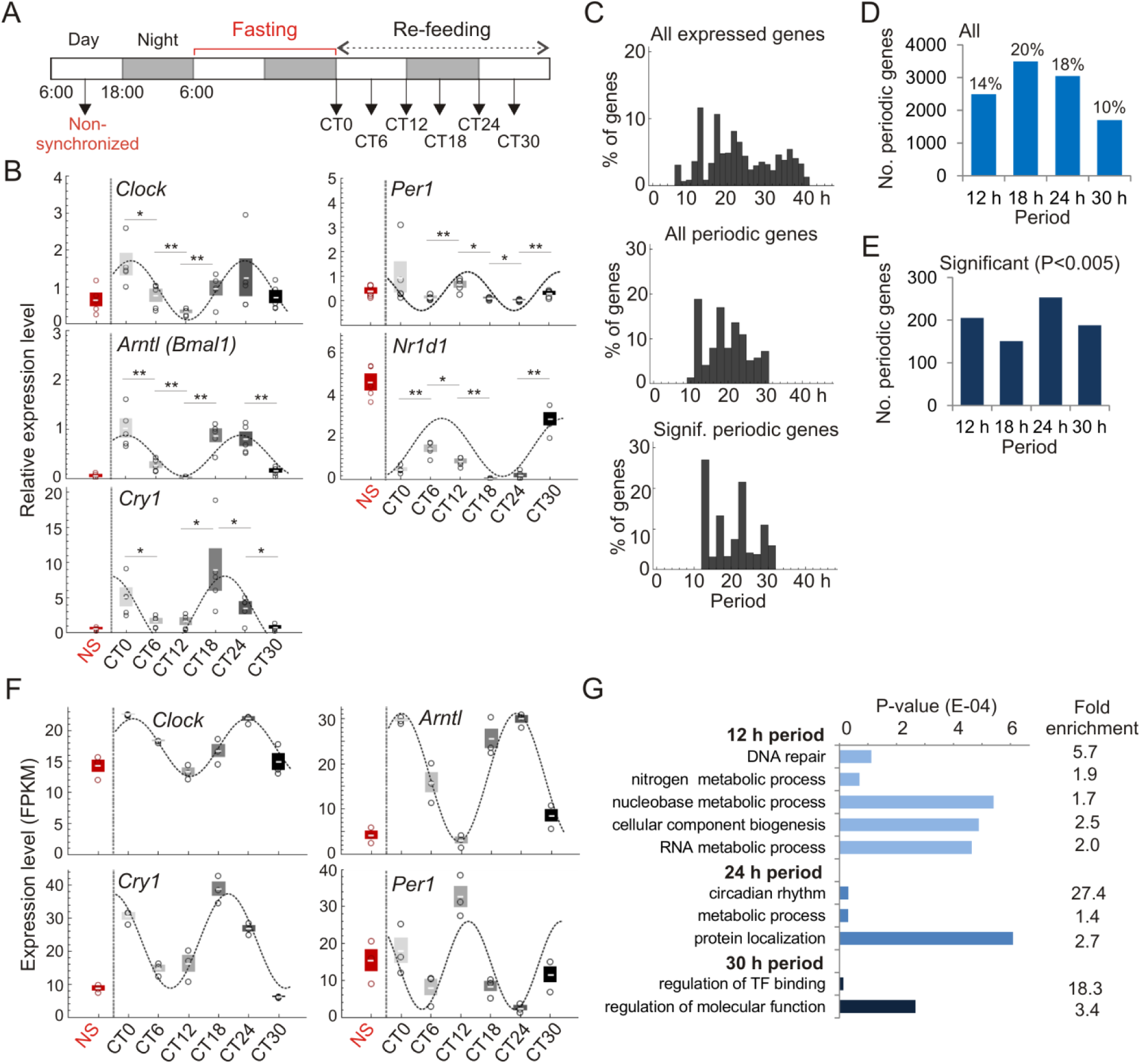
Fasting and refeeding entrains circadian liver gene expression. (A) Circadian clock entrainment schedule used in this study. Mice were fasted for 24 h and refed at 6:00 am (CT0; light on, white bar; black bar, light off). Livers were collected at each CT (h) and from non-synchronized mice. (B) RT-qPCR analysis of expression of central clock genes in non-synchronized (NS) mice and from CT0 to CT30 (mean ± SD, single data points; 3 mice at NS and per CT). Cosine curves are best-fits determined by MetaCycle. (C) Period distribution for all 17,330 expressed genes in the liver (top), all periodic genes (middle) and significantly (*P* < 0.005; Fisher’s exact test) periodic genes (bottom). (D) Numbers of genes oscillating with indicated periods; % of all expressed genes are also shown. (E) Numbers of significantly periodic genes (*P* < 0.005; Fisher’s exact test) for each period. (F) Circadian expression of central clock-control genes (RNA-seq FPKM; 3 mice per CT; mean ± SD, individual data points); cosine curves are MetaCycle best-fits. (G) Enriched GO terms for periodic genes. See Supplemental Tables S1 and S2 for gene lists and GO terms, respectively. No significant GO term enrichment was found in the 18 h period.

We used MetaCycle (Wu et al. 2016) to identify genes with periodic expression patterns from liver RNA-sequencing (RNA-seq) data generated at each CT. Assessing period the distribution for all 17,330 expressed genes (i.e. expressed at at least one time point), we found dominant oscillations at 12, 18, 24 or 30 h (Fig. 1C, top panel); thus we focused our analyses to oscillations with 12, 18, 24 and 30 h periods, each ± 3 h, i.e. half the time-resolution in our study (Fig. 1C, middle panel). Nearly 20% of oscillations are circadian (24 h period; 3,046 genes), and thousands genes oscillate with periods within the circadian rhythm (12 h: 2,496 genes; 18 h: 3,495 genes) and beyond (30 h: 1,702 genes; Fig. 1D), in line with previous reports (Hughes et al. 2009; Korencic et al. 2014; Zhu et al. 2017). Among those, 151-253 display significantly oscillating transcripts at the *P* < 0.005 level (Fisher’s exact tests; Fig. 1C, lower panel; Fig. 1E). We also find 644 genes with significant (*P* < 0.005) mRNA oscillations with a period outside 12, 18, 24 or 30 h (see **Supplemental Table S1** for lists of periodic genes). We also defined a set of ‘non-periodic’ genes based on a P-value of 1 > *P* > 0.9999…. with variations in transcript levels discordant with any cosine curve. Periodic genes included the core clock genes (Fig. 1F; **Supplemental Fig. S1A**), corroborating entrainment of the clock.

Gene ontology analysis confirms enrichment of 24 h periodic genes in rhythmic and circadian processes, including key metabolic functions (Fig. 1G; **Supplemental Table S2**). In addition, 12 h periodic genes are mainly enriched in DNA repair functions (Fig. 1G). These include *Fan1*, encoding a DNA repair nuclease, the E3-ubiquitin ligase-encoding gene *Rad18*, the ATP-dependent DNA helicase-encoding gene *Recql4*, the SWI/SNF subunit-coding gene *Smarcb1*, and the poly(ADP-ribose) glycohydrolase-encoding gene *Parg* (**Supplemental Table S1**). The RNA demethylase-encoding gene *Alkbh5* also displays 12 h oscillations, raising the hypothesis that RNA modification may occur in a periodic manner. *Med1*, which encodes the DNA looping factor and mediator component MED1, also exhibits a 12 h period, in line with the circadian binding profile of MED1 to chromatin (Kim et al. 2018). 30 h periodic genes are primarily involved in the regulation of TF binding (Fig. 1G); these include *Ppargc1b* and the CREB-regulated transcription co-activator *Crtc3*. These results illustrate the involvement of periodic genes in important chromatin repair and regulation functions.

For a given period, we defined mRNA oscillations in phase and in opposition of phase, i.e. with respectively maxima and minima occurring at the start of the periodic cycle (Fig. 2A). Given the 6 h time-resolution in our study, we find that 12 h oscillations occur in phase or in opposition of phase, while 18, 24 or 30 h oscillations can in addition be offset by one-quarter cycle (Fig. 2A,B; **Supplemental Fig. S1B,C**). Interestingly, a number of periodic genes encode BTB/POZ domain TFs, some of which are involved in targeting chromatin to the nuclear lamina(Zullo et al. 2012) (**Supplemental Table S3**). Some of these genes oscillate in opposition of phase and are CLOCK or BMAL1 targets (e.g. *Zbtb40, Zfp608*; Fig. 2C), putatively linking periodic expression of these BTB/POZ domain TFs with periodic associations of chromatin with the nuclear envelope.

**Figure 2.**
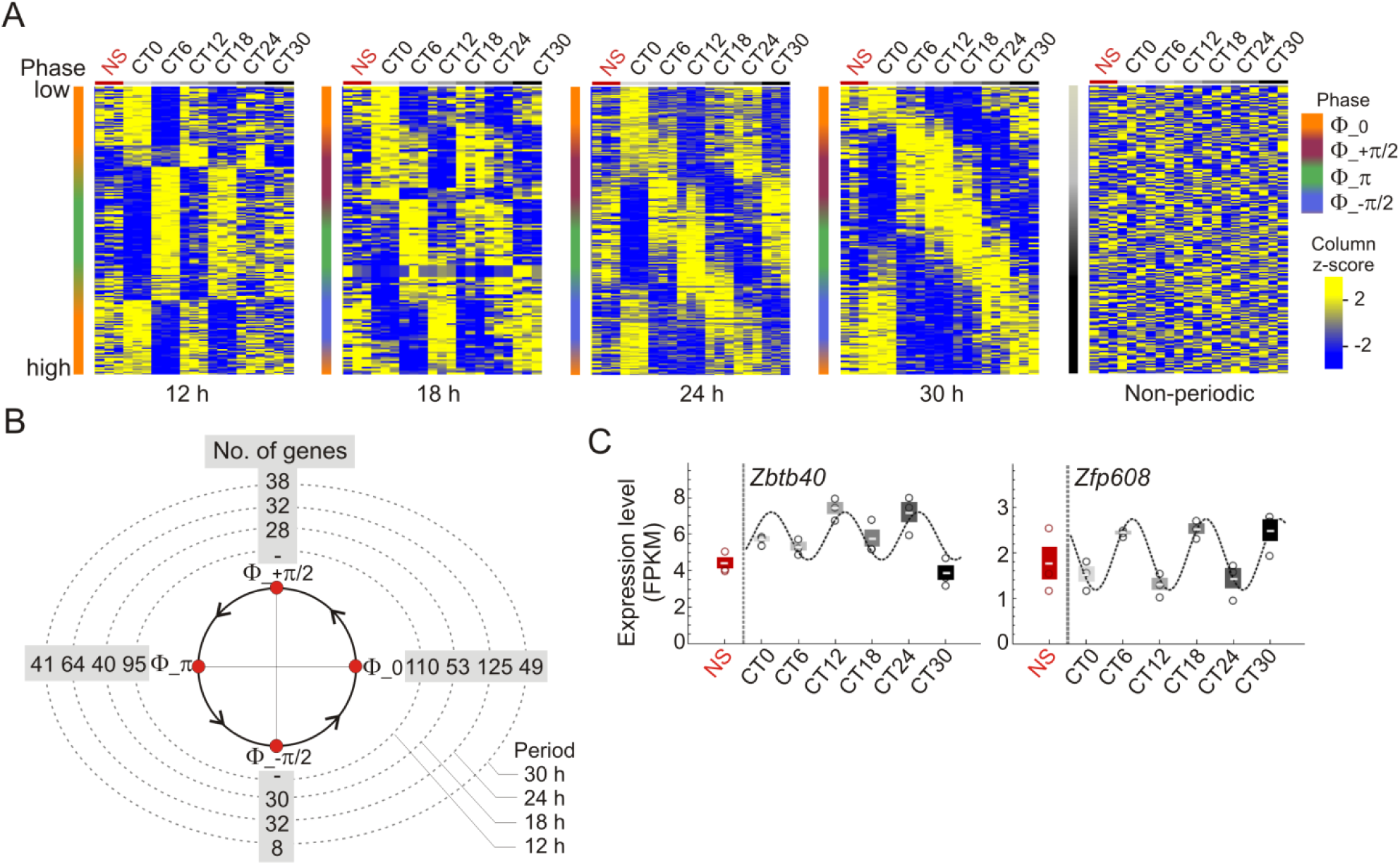
Periodic genes oscillate with distinct phases. (A) Expression profiles (FPKM z-scores) of periodic and non-periodic genes in NS mice and from CT0 to CT30 (3 mice per CT). Genes are ranked by increasing phase value along the y axis (see scale). (B) Trigonometric clock and number of significantly (*P* < 0.005; Fisher’s exact test) periodic genes with indicated period and phase. (C) Expression profiles of the BTB/POZ domain TF gene *Zbtb40* and the zinc-finger TF gene *Zfp608* (RNA-seq FPKM; 3 mice per CT; mean ± SD, individual data points); cosine curves are MetaCycle best-fits.

### Entrainment of the circadian clock resets LMNB1-chromatin interactions

We therefore next examined chromatin association with the nuclear lamina during the circadian cycle. First, we show that LMNB1 protein and *Lmnb1* transcripts are expressed without significant variations in liver (Fig. 3A,B; **Supplemental Fig. S1D**). Second, chromatin immunoprecipitation (ChIP)-qPCR of LMNB1 from liver confirms LMNB1 enrichment in established mouse constitutive (c)LADs (Meuleman et al. 2013) (**Supplemental Fig. S2A,B**), and validates LAD variations detected by ChIP-sequencing (ChIP-seq) of LMNB1 at three distinct time points (**Supplemental Fig. S2C,D**).

**Figure 3.**
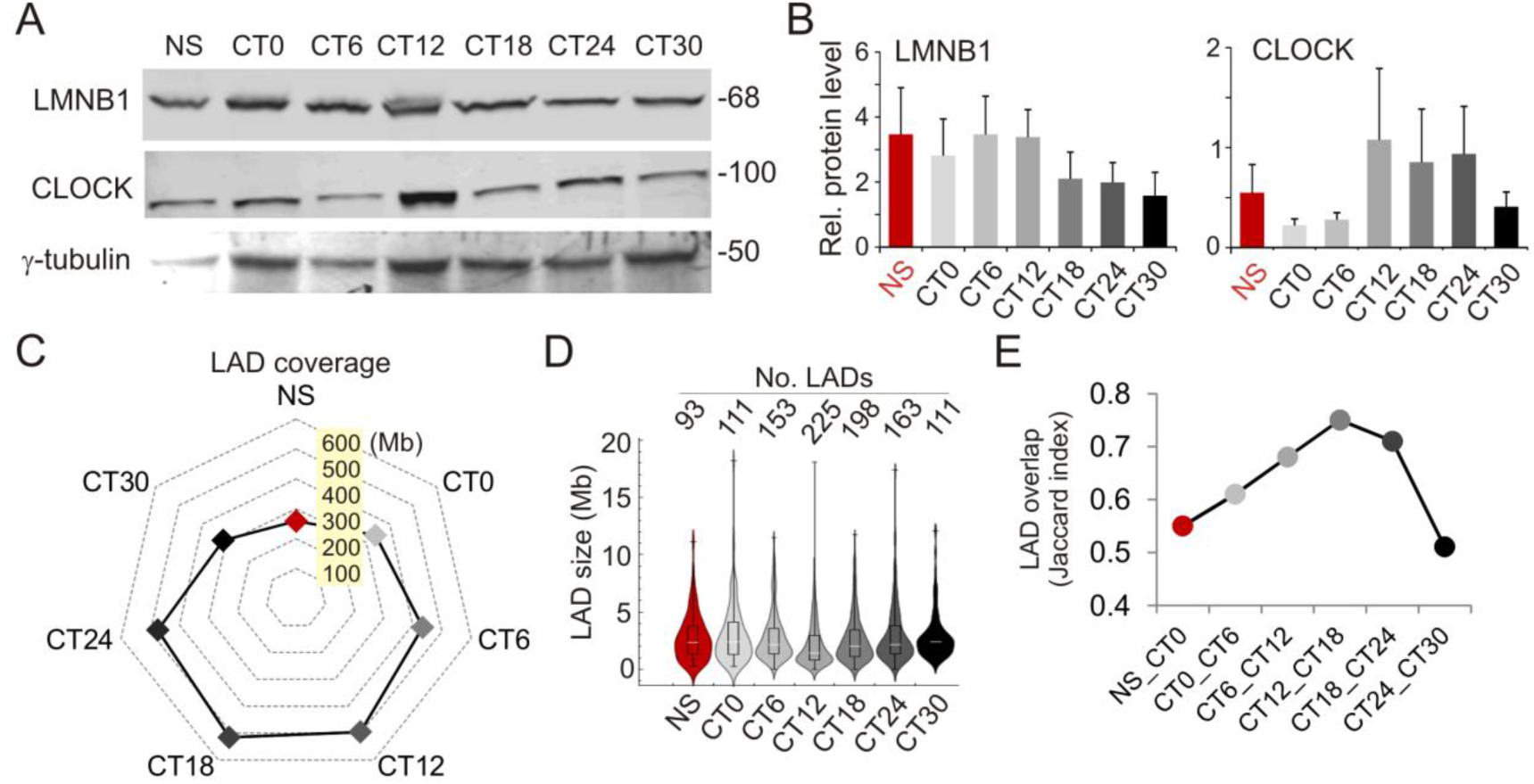
Characterization of LADs during the circadian cycle. (A) Western blot of lamin B1 (LMNB1) and CLOCK; γ-tubulin was used as loading control. (B) Quantifications of immunoblots as in (A) relative to γ-tubulin; mean ± SD from analysis of blots from livers of 3 mice per CT. (C) Radar plot representation of genome coverage by LADs during the circadian cycle (values, in Mb, for intersects of LAD coverage between replicates); circumference scale range is 0-600 Mb. (D) Distribution and median of LAD sizes at each CT (LAD intersects at each CT); see Supplemental Fig. S3A for all replicates. (E) Jaccard indices of LAD overlap between consecutive CTs (LAD intersects at each CT); see also Supplemental Fig. S3C.

Thus, we determined by LMNB1 ChIP-seq to what extent LADs varied during the circadian cycle. LAD sizes are overall stable and display low gene density at all time points (**Supplemental Fig. S3A,B**). Analysis of between-replicate LAD intersects reveals an increase in LAD coverage from CT0 (339 Mb) to CT18 (516 Mb), followed by a decrease to CT30 (312 Mb) (Fig. 3C); this primarily results from variations in LAD numbers rather than size (Fig. 3D). Nevertheless, LADs display a marked overlap across replicates and CTs, yet with the next-to-lowest overlap between non-synchronized (NS) mice and mice at CT0 (Fig. 3E**; Supplemental Fig. S3C**). Lowest LAD overlap was detected between CT24 and CT30 (Fig. 3E), in line with a view of a loss of synchrony as time progresses and as reflected by the transcriptome (see Fig. 2A). These results reveal the overall conservation of LADs in mouse liver during the circadian cycle; however they also indicate that variable LMNB1-chromatin interactions do occur.

We examined variable (v)LADs more closely and first assessed to what extent fasting/refeeding reset LADs. LADs from non-synchronized (NS) mice and CT0 LADs reveal substantial differences in coverage (Fig. 4A,B), which was anticipated from the Jaccard index (0.55; see Fig. 3E). Entrainment of the clock is manifested by a gain of 126.7 Mb at CT0, of which 79.7 Mb are stand-alone LADs and 47 Mb represent extensions of pre-existing LADs (Fig. 4B). LMNB1 enrichments at CT0 were confirmed by ChIP-qPCR (Fig. 4C,D**; Supplemental Fig. S3D**). There is also a loss of 46 Mb at CT0 mostly as stand-alone LADs (Fig. 4B). Thus entrainment of the clock is associated with a resetting of LADs manifested by a net gain in LMNB1-chromatin interactions.

**Figure 4.**
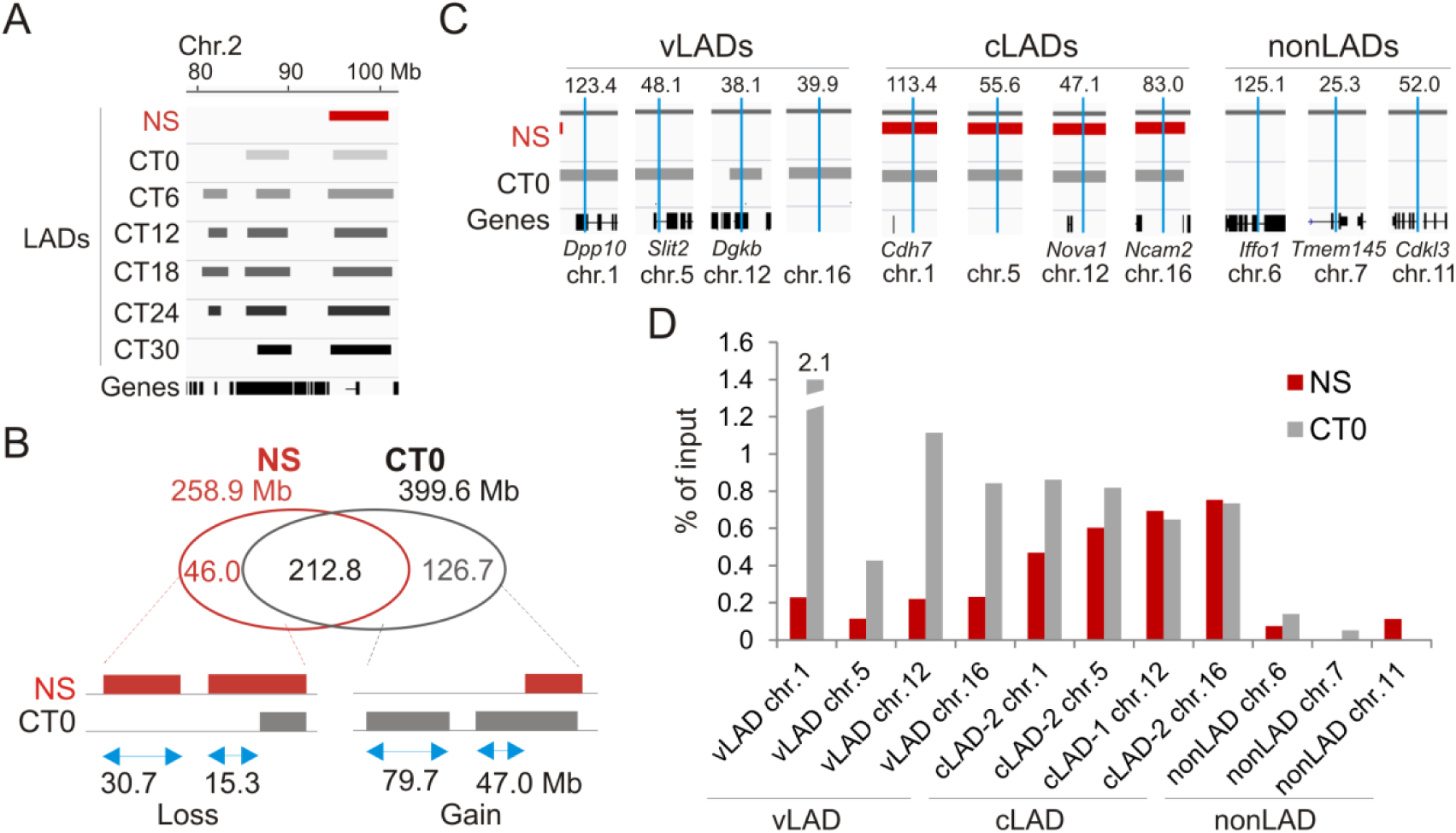
Resetting of LADs after entrainment of the circadian clock. (A) Browser view of LADs in a region of chromosome 2; LAD intersects between replicates at each CT are shown. (B) Venn diagram analysis of LAD overlap between NS mice and CT0 (intersect of replicates); bottom, genome coverage by LADs gained or lost at CT0 relative to NS, for stand-alone LADs and LAD extensions or shortenings (values for LAD intersects at NS and CT0). (C) Browser views of vLAD, cLAD and nonLAD regions, with position of amplicons analyzed by ChIP-qPCR in (D) (blue bars). (D) ChIP-qPCR analysis of LMNB1 enrichment in indicated vLAD, cLAD and nonLAD regions; see Supplemental Table S7 for position of amplicons.

### Periodic oscillations in LMNB1-chromatin interactions constitute a minor fraction of variable LADs

We next examined vLADs more closely during the circadian cycle. vLADs occur by LAD extension or shortening, sometimes resulting in a fusion or splitting of LADs, or by formation and dissociation of entire LADs (see Fig. 4A). We therefore devised a strategy to quantitatively characterize the genomic coverage of these vLADs over time. We determined, for each LAD, the maximal coverage (cov_max_) of the LAD in the CT0-CT30 time-course, as the union of LAD coverage across all biological replicates and CTs, and ascribed to this cov_max_ a reference value of 1 (Fig. 5A). The cov_max_ 5’ and 3’ boundaries provided genomic coordinates for measures of variations in LAD length within this area. For each CT and replicate, variable 5’ and 3’ lengths (in base pairs) were extracted and standardized from 0 to 1 (cov_max_), 0 referring to the complete disappearance of a LAD. We then applied the MetaCycle tool to identify any periodicity in the extension or shortening of LADs within the cov_max_ area (Fig. 5A).

**Figure 5.**
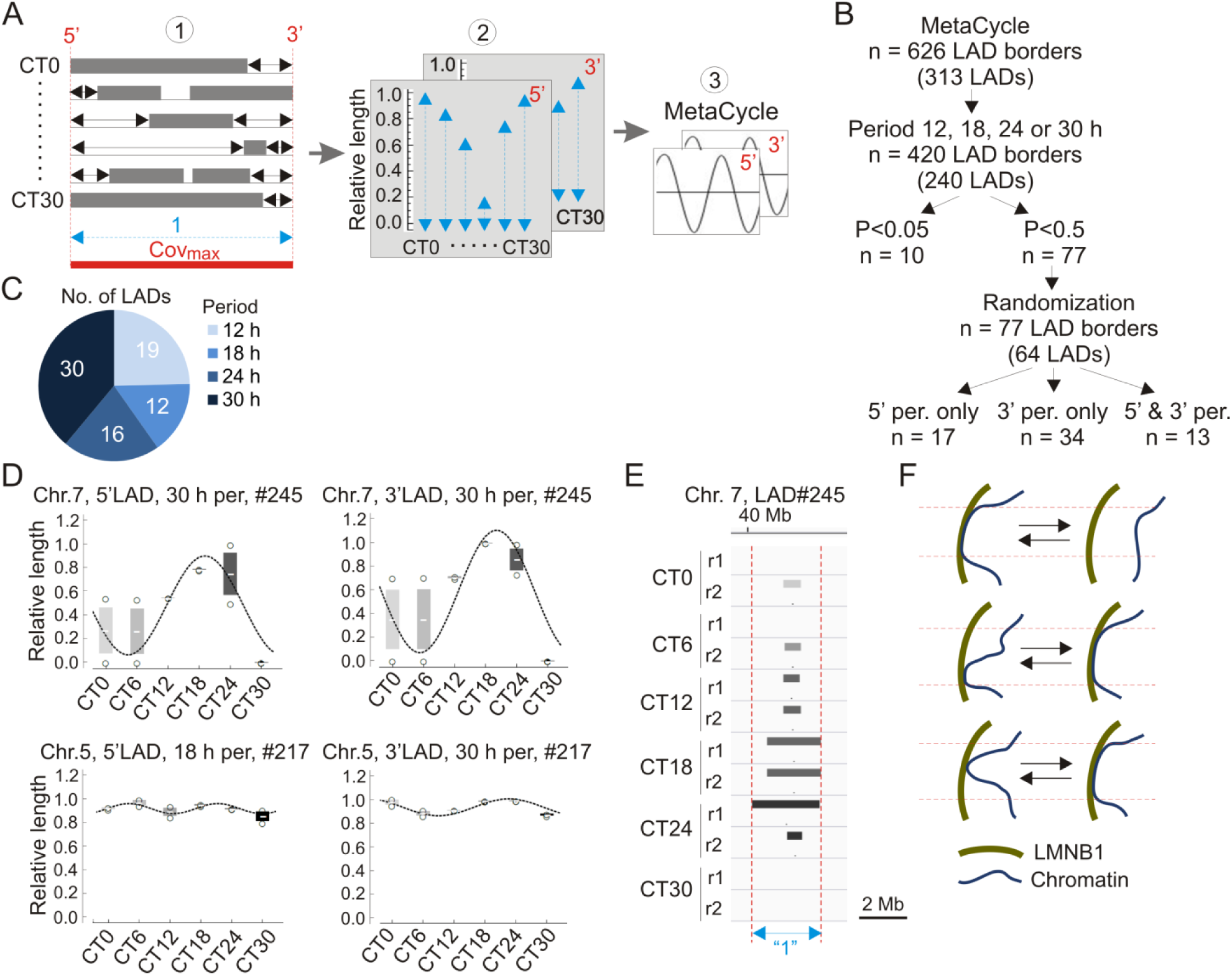
Identification of periodic LADs. (A) Approach to the identification of periodic LADs. (1) A variable LAD area is identified across all CTs and replicates; the maximal merged area of these LADs is defined (Cov_max_; red bar) and distances from the 5’ and 3’ end of each LAD to Cov_max_ limits are determined (black arrows); (2) relative lengths are calculated on both ends (1 = Cov_max_ length; 0 = no LAD); (3) MetaCycle is applied to identify periods at the 5’ and 3’ ends of LADs. (B) Identification of periodic LADs using MetaCycle. (C) Period distribution of the 77 oscillating LAD borders. (D) Examples of periodic oscillations in LAD 5’ and 3’ length during the circadian cycle (see Supplemental Table 5 for LAD features). Mean ± SD, individual data points and MetaCycle best-fit cosine curves are shown. (E) Browser view of a periodic LAD (LAD #245; see also panel D). (F) Summary of oscillatory LAD patterns after entrainment of the clock. Arrows refer to transitions between consecutive time points.

We identify 626 vLAD borders (313 LADs) with oscillating lengths over our experimental time-frame (Fig. 5B). We then identify 12, 18, 24 or 30 h (± 3 h) 5’ and/or 3’ periodic oscillations in 420 of these LAD borders (in 240 LADs). Ascribing a *P* value of < 0.05 (Fisher’s exact test) yields 10 significantly periodic LADs with periods evenly distributed throughout the circadian cycle (**Supplemental Table S5A**). Relaxing the *P* value to *P* < 0.5 logically increases the output number of periodic LAD borders to 77 (30 5’-periodic and 47 3’-periodic) among 64 distinct LADs (Fig. 5B,C; **Supplemental Table S5A**). These periodic LADs withstand a randomization test of all replicates and time points (Fig. 5B), suggesting that LAD periodicity observed in our data is imposed by the order of CTs. Among these 64 periodic LADs, 17 display 5’-only oscillations, 34 show 3’-only oscillations, and 13 display both 5’ and 3’ oscillations (Fig. 5B,D; **Supplemental Table S5B**), including 7 with identical 5’ and 3’ periods (Table 1; Fig. 5D). Periodic LADs include LADs with a constitutive core and variable borders, and LADs that entirely appear or disappear (Fig. 5D,E).

**Table 1.** Period of 5’ and 3’ LAD borders when both periodically oscillate.

| LAD No. | Chr. | 5' period (h) | 3' period (h) |
| --- | --- | --- | --- |
| 37 | 10 | 30 | 30 |
| 56 | 12 | 12 | 24 |
| 78 | 14 | 18 | 12 |
| 124 | 17 | 18 | 24 |
| 160 | 2 | 12 | 12 |
| 201 | 4 | 30 | 30 |
| 211 | 5 | 30 | 30 |
| 214 | 5 | 12 | 18 |
| 217 | 5 | 18 | 30 |
| 245 | 7 | 30 | 30 |
| 262 | 8 | 18 | 12 |
| 287 | X | 12 | 12 |
| 288 | X | 24 | 24 |

See Supplemental Table S5 for LAD features.

We conclude that the majority of LMNB1-chromatin interactions are conserved during the circadian cycle, but these are also accompanied by discrete oscillations in such interactions. Nevertheless, significant periodicity in 5’ or 3’ LAD border interactions with LMNB1 only applies to a minority of vLADs with overall little synchrony between borders.

### Periodic gene expression is uncoupled from rhythmic LMNB1-chromatin interactions

We then assessed the relationship between periodic gene expression and periodic LADs. The vast majority of significantly periodic genes are outside LADs at any time point during the circadian cycle, with 0-2% of periodic genes found in periodic LADs (Fig. 6A). Among these, *Ppargc1a* (18 h period; *P* = 0.00055), encodes the PPARG co-activator PGC1A, a regulator of neoglucogenesis in liver through activation of *Foxo1*, which is also periodic, but not in a LAD. This raises the hypothesis that metabolic functions may centrally be under influence of periodic interaction of genes with LMNB1; however our data argue that is not a predominant phenomenon.

**Figure 6.**
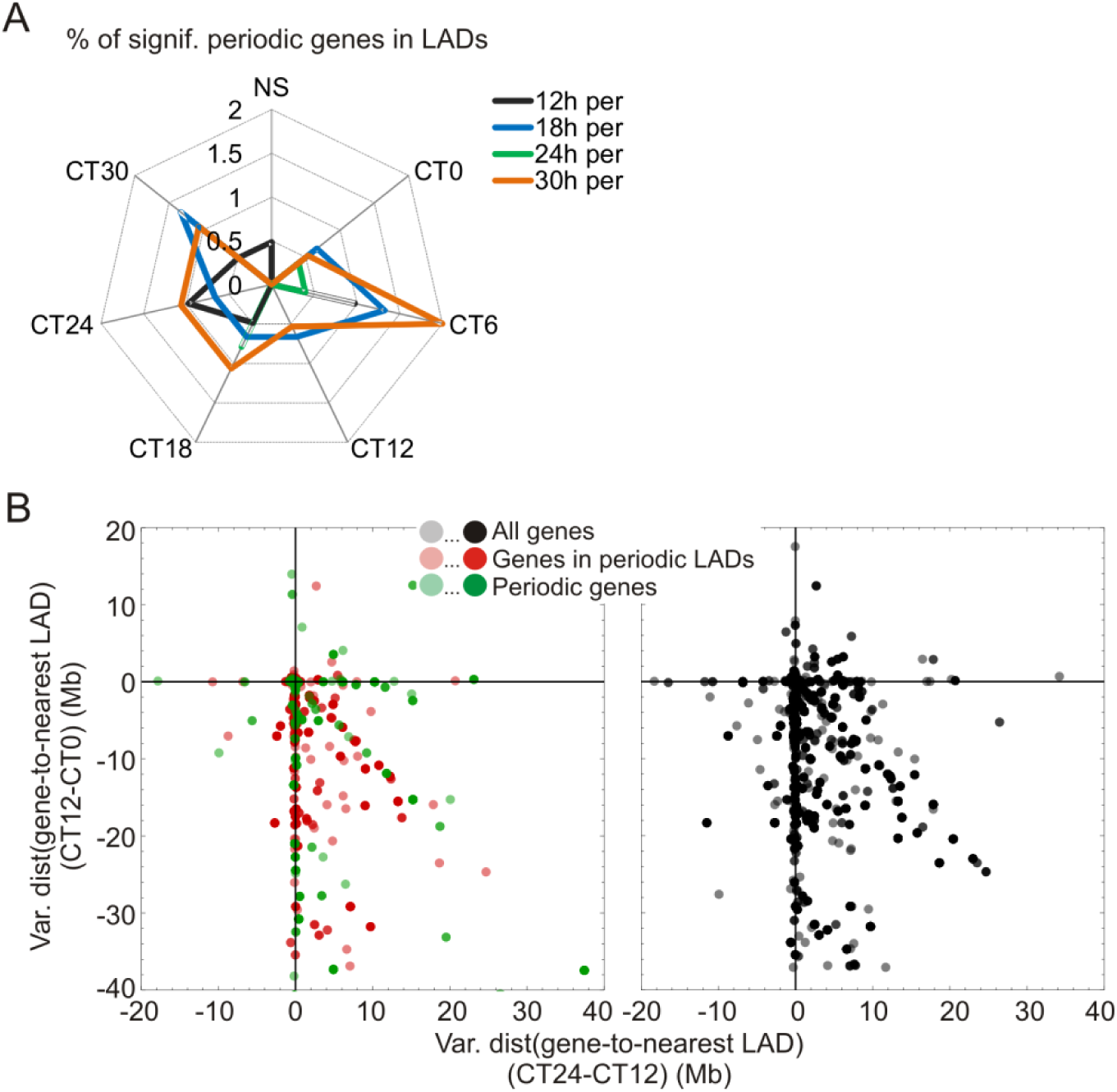
Periodic LADs and gene-LAD distances are uncoupled from periodic gene expression. (A) Percentages of significantly periodic genes (*P* < 0.005, Fisher’s exact test) found in LADs. (B) Scatter plot of variations in (gene-to-nearest LAD) distance between CT0 and CT12 (y axis) and between CT12 and CT24 (x axis). Data are shown for all genes (gray scale, right plot) and for all periodic genes (green scale, left) genes in periodic LADs (red scale, left). Color intensity of data points reflects the number of genes at a particular coordinate on the graph. All genes are shown separately (right) for clarity.

Since most genes reside outside LADs during the circadian cycle (LADs are gene-poor; see **Supplemental Fig. S3B**), we next determined the relationship between periodic genes and gene distance to the nearest LAD. Consistent with the variation in LAD coverage seen during the circadian cycle and the gene-poor nature of LADs, we observe an overall decrease in gene-nearest LAD distance between CT0 and CT12, and an increase thereafter (Fig. 6B). However, there is no correlation between gene periodicity and variation in gene-to-nearest LAD distance, and this regardless of the magnitude of variations in gene-LAD distances during the circadian cycle (Fig. 6B, green data points). Similarly, central clock-regulating genes are typically megabases away from the nearest LAD (**Supplemental Fig. S4A**; **Supplemental Table S4**), and transcription start sites and promoter regions of these genes are essentially devoid of LMNB1 at any CT (**Supplemental Fig. S4B**). This altogether suggests the maintenance of a lamin-less environment permissive for transcriptional activation during the circadian cycle. Our findings therefore argue that oscillatory expression of periodic genes, including central clock-control genes, is uncoupled from a direct association with the nuclear lamina or from their localization in the vicinity of variable LADs.

## Discussion

Oscillations in chromatin conformation mediated by rhythmic chromosomal interactions contribute to the regulation of circadian gene expression (Aguilar-Arnal et al. 2013; Xu et al. 2016; Kim et al. 2018; Mermet et al. 2018; Yeung et al. 2018). Here, we applied MetaCycle, a tool designed to identify periodic transcript oscillations (Wu et al. 2016), to identify periodic changes in genome coverage by LMNB1. Our approach distinguishes 5’ and 3’ LAD extension or shortening from entire LAD emergence or disappearance (Fig. 5F). We provide evidence of a net gain of lamin-chromatin interactions by entrainment of the circadian clock, which speculatively may reinforce the robustness of segregation of chromatin domains. This is followed by periodic interactions of specific regions with LMNB1 suggestive of discrete rhythmic associations with the nuclear lamina. Our data add to mounting evidence that chromatin is able to display oscillations in spatial conformation (Kim et al. 2018).

Given the overall repressive chromatin environment at the nuclear periphery (van Steensel and Belmont 2017), and evidence of cyclic recruitment and silencing of specific genes at the nuclear envelope in a human cancer cell line (Zhao et al. 2015), we reasoned that periodic gene expression could at least in part be regulated by periodic interactions with the nuclear lamina. Counterintuitively however, several elements in our data argue that in mouse liver, periodic gene expression is uncoupled from, and thus unlikely to be regulated by, periodic chromatin associations with the nuclear lamina. Genes with periodic transcript levels reside outside LADs at any time point during the circadian cycle; this includes *Pard3*, which has been found to be periodically recruited to the nuclear envelope in human colon cancer cells (Zhao et al. 2015), but in mouse liver is localized 15 Mb away from a conserved LAD. Similarly, central clock-control genes reside megabases away from the nearest LAD and show no promoter association with LMNB1. This configuration may keep these genes in a transcriptionally permissive environment throughout the circadian cycle, compatible with a regulation of circadian transcription by rhythmic TF binding and activity (Koike et al. 2012; Masri et al. 2014; Zhang et al. 2015). Periodic genes can also flank constitutive LADs and conversely, genes with stable (high or low) expression levels may flank oscillatory LADs. Lastly, we find no relationship between periodic gene expression and gene-nearest LAD distance during the circadian cycle. Thus, the periodic LAD patterns identified here in liver cannot readily explain oscillatory gene expression. Rather, periodic hepatic gene expression appears to be uncoupled from periodic chromatin association with the nuclear lamina.

Concurring with our results, other chromatin-linked processes are uncoupled from gene expression patterns. For example, many oscillatory genes display stable chromosomal interactions (Kim et al. 2018), including promoter-enhancer contacts (Beytebiere et al. 2018), during the circadian cycle. Conversely, most expressed genes harbor circadian promoter and enhancer histone modifications irrespective of transcriptional periodicity (Koike et al. 2012). These observations highlight a complex cross-talk between circadian transcription and rhythmic changes in chromatin states.

Our findings raise several important issues. First, periodic LMNB1-chromatin interactions are not a prominent feature of variable LADs during the circadian cycle. Even though hundreds of LADs display oscillatory variations in 5’ and 3’ end coverage, we only find with our approach a handful of significantly periodic LADs (13 in our dataset), with half displaying synchronous oscillations between their 5’ and 3’ borders. A lenient *P* value increases the number of ‘periodic’ LADs, for which our data suggest that rhythmicity is elicited by the underlying order of CTs (i.e. circadian time). Nevertheless, these LAD oscillations are again largely asynchronous between 5’ and 3’ borders. Our findings suggest therefore that periodicity in lamin-chromatin interactions identified here may reflect stochasticity among variable LADs during the circadian cycle in the liver, or involve a level of regulation that currently remains unappreciated.

Second, how would periodic lamin-chromatin interactions, then, be regulated? The cistrome of circadian genes can oscillate in a manner concordant with circadian gene expression (Feng et al. 2011; Fang et al. 2014; Zhang et al. 2015), thus factors mediating chromatin-lamin interactions may also be periodically recruited to target loci. Several genes encoding BTB/POZ domain proteins oscillate during the circadian cycle. These proteins share DNA binding motifs enriched in LADs (Guelen et al. 2008) and in lamin-associated sequences (Zullo et al. 2012; Lund et al. 2013), and are found in sequences able to re-localize chromatin to the nuclear lamina (Harr et al. 2015). Thus, oscillating LADs could potentially be regulated through periodic recruitment of factors important for chromatin localization at the nuclear periphery (Shachar et al. 2015).

Third, what is the significance of oscillatory chromatin-lamin interactions and their impact on genome architecture? Resetting of LADs after entrainment of the clock reflects a synchronization process leading to a reduction in cell-to-cell variation in LAD profiles at the organ level. Robust LAD ‘marking’ may strengthen the robustness of liver-specific gene expression, possibly by segregation of heterochromatin from euchromatin. Oscillations of subsets of LADs, regardless of periodicity, may establish local and dynamic radial chromatin states in regions that are in 3-dimensional proximity, but not necessarily in linear proximity, affecting gene expression in these regions (Paulsen et al. 2019). Thus periodic LAD displacement may result in radial repositioning of topological chromatin domains (Robson et al. 2016). Assessment of periodic LADs in a 3-dimensional environment, even linearly away from LADs, should shed light on the putative long-range *cis* or *trans* impact of circadian LAD dynamics on genome architecture and function.

## Methods

### Mice

Wild-type C57Bl/6 male mice (Jackson Laboratories) were housed in 12 h light/12 h dark cycles with lights on at 6 a.m. and lights off at 6 p.m. Mice were kept off chow for 24 h, refed *ad libitum* at circadian time CT0 (6 a.m.) and sacrificed at CT6, 12, 18, 24 and 30 h (n = 7 mice per CT). Non-synchronized mice (n = 7) were sacrificed at 12:00 noon on the day prior to food restriction. Livers were collected from all mice, partitioned and snap-frozen in liquid nitrogen. Procedures were approved by the University of Oslo and Norwegian Regulatory Authorities (approval No. 8565).

### RNA-seq and gene expression analysis

Total RNA was isolated from livers of five mice at each CT using the RNeasy Mini Kit (Qiagen). TNA-seq libraries (n = 3 mice per CT) were sequenced on an Illumina HiSeq2500. RNA-seq reads were processed with Tuxedo (Trapnell et al. 2010). TopHat v2.10 was used to align reads with no mismatch against the mm10 genome applying the Bowtie2 preset option ‘b2-very sensitive’ (Langmead and Salzberg 2012; Kim et al. 2013). Transcript level was estimated using cufflinks v2.2.1, and differential gene expression determined using cuffdiff v2.2.1 (Trapnell et al. 2010). Expression plots display mean ± SD relative level (RT-qPCR; n= 5 mice) or FPKM (n = 3 mice) per CT, with single data points. A gene was ascribed to a LAD if its transcription start site overlapped with the LAD.

### Metacycle analysis

We used MetaCycle (Wu et al. 2016) to identify genes with periodic expression patterns. MetaCycle measures the goodness-of-fit between RNA-seq FPKM and theoretical cosine curves with varying periods and phases. The extrapolated periodic function best fitting the RNA-seq data is selected and significance of a given periodicity is determined by assigning *P* values after scrambling FPKM values. MetaCycle was applied to the entire range of CTs. To fit RNA-seq data with periodic functions, Metacycle normalizes FPKM values by computing z-scores. Our time-series data are integer intervals with even sampling and do not include missing values (3 replicates RNA-seq data; 2 replicates LMNB1 ChIP-seq data per CT; see also Determination of oscillating LADs). Given the features of our time-series data, MetaCycle incorporated both the JTK_CYCLE (JTK) and the Lomb-Scargle (LS) methods for periodic signal detection. Based on our data and time-resolution, available period values are real numbers ranging from 12 to 48 h for JTK, and integers (0, 6, 12, 18, 24, 30 or 36 h) for LS. Periods reported in our analyses are the mean of JTK and LS period values. Of note, the 6-h resolution time-course in our study did not allow detecting changes in gene expression periods under 12 h. Moreover, the restricted 12 ± 3 h (i.e. half the time-resolution in our study) period group did not able us to distinguish groups of 12 h periodic genes oscillating in positive (ϕ_π/2) or negative (ϕ_-π/2) quadrature of phase, but only those oscillating in phase or opposition of phase.

### Immunoblotting

Proteins were separated by 10% SDS-PAGE, transferred onto an Immobilon-FL membrane (Millipore) and membranes blocked with Odyssey blocking buffer (LI-COR). Membranes were incubated with antibodies against CLOCK (1:500; Abcam ab3517), LMNB1 (1:1000; Santa Cruz Biotechnology sc6216) and γ-tubulin (1:10000; Sigma-Aldrich T5326). Proteins were detected with IRDye-800- or IRD IRDye-680-coupled secondary antibodies.

### ChIP

ChIP of LMNB1 was done as described (Rønningen et al. 2015) and adapted for liver pieces. Snap-frozen liver tissue pieces (40-50 mg) were thawed on ice and minced on ice for 30 sec. Minced tissue was resuspended in PBS with 1 mM PMSF and protease inhibitors, and homogenized by 7-10 strokes in a 2-ml Dounce homogenizer using pestle B (tight-fitting). Samples were centrifuged at 400 g and supernatants discarded. Pellets were resuspended in PBS containing 1% formaldehyde and cross-linking was allowed to occur for 10 min at room temperature. Cross-linked samples were sedimented and lysed in RIPA buffer (140 mM NaCl, 10 mM Tris-HCl, pH 8.0, 1 mM EDTA, 0.5 mM EGTA, 1% Triton X-100, 1% SDS, 0.1% sodium deoxycholate, 1 mM PMSF, protease inhibitors). To generate 200-500 bp DNA fragments, cells were sonicated 4 times 10 min in a Bioruptor (Diagenode). After sedimentation, the supernatant was diluted 10-fold in RIPA and incubated with anti-LMNB1 antibodies (10 µg; Abcam ab16048) coupled to Dynabeads Protein G (Invitrogen) overnight at 4°C. ChIP samples were washed 3 times in ice-cold RIPA. Cross-links were reversed and DNA eluted for 6 h at 68°C in elution buffer (50 mM NaCl, 20 mM Tris-HCl, pH 7.5, 5 mM EDTA, 1% SDS) containing 40 ng/ml proteinase K. DNA was extracted and libraries prepared and sequenced on an Illumina HiSeq2500.

### ChiP-seq data processing

LMNB1 ChIP-seq and input reads were mapped to the mm10 genome using Bowtie v2.25.0 (Langmead et al. 2009) with default parameters after removing duplicates using Picard’s MarkDuplicates. To avoid normalization bias, we ensured that each pair of mapped ChIP and input read files had the same read depth by down-sampling reads for each chromosome individually. Mapped reads were used to call LADs using Enriched Domain Detector (Lund et al. 2014) with the following alteration. To account for technical variation potentially occurring in LAD calling, we first ran EDD 10 times on each lamin ChIP-seq dataset in auto-estimation mode for GapPenalty (GP) and BinSize (BS). Average GP standard deviation was ≤ 1.6 units while BS did not vary. GP variations elicited minimal alterations in LAD calls, allowing estimation of technical variability. For all LADs, median length of these variations was 0.32 Mb. This represents < 1% of total LAD coverage, and is 3-15 times smaller than median LAD sizes for each CT and replicate. Thus, intrinsic EDD variability did not significantly impact LAD calling. Average GP and BS values from the 10 runs were used to set GP and BS before a final EDD run with each LMNB1 ChIP-seq dataset (**Supplemental Table S8**). Intersects between LADs and genes were determined using BEDTools v2.21.0 (Quinlan and Hall 2010) and BEDOPS v2.4.27 (Neph et al. 2012). Scripts were written in Bash, Perl (Stajich et al. 2002) or R (Team 2015) and ggplot2 in R was used for plots. Browser files were generated by calculating ChIP/input ratios in 10-kb bins with input normalized to the ratio of total ChIP reads over total input reads.

### Determination of gene-LAD distance

Gene-to-nearest LAD distance was calculated as the distance from the 5’ and 3’ end of a gene to, respectively, the nearest 3’ and nearest 5’ LAD border. Gene strand was respected in the calculation and LAD intersects were used at each CT. If a gene was entirely within a LAD, gene-to-nearest LAD distance was the distance from the 5’ and 3’ end of the gene to the first neighboring LAD, both upstream and downstream. Moreover, in our approach, the internal configuration of a variable LAD within the cov_max_ area (e.g. a LAD split or fusion) is not taken into account in the determination of periodicity in the 5’ and 3’ borders.

### Determination of periodic LADs

The approach is outlined in Figure 5A and described in the results. In brief, to quantify the genomic coverage of variable LADs, we determined for each LAD its ‘maximal coverage’ (cov_max_) as the union of LAD coverage across all biological replicates and CTs; we ascribed to this cov_max_ a relative reference value of 1. The cov_max_ 5’ and 3’ borders provided genomic coordinates for measures of variations in LAD length within the cov_max_ area. For each CT and replicate, variable 5’ and 3’ genomic lengths (in base pairs) were extracted and standardized from 0 to 1, 0 referring to the complete disappearance of a LAD and 1 being the cov_max_ value. MetaCycle was then applied to determine the periodicity of LAD extensions and shortenings at 5’ and 3’ borders.

### Data viewing

Browser views were produced with the Integrated Genomics Viewer (Robinson et al. 2011) using mm10 genome annotations.

### Statistics

FPKM or RT-qPCR transcript levels were compared using unpaired t-tests. Metacycle used its built-in statistical method to assign *P* values from Fisher’s exact tests. The JTK and LS methods used in Metacycle assign a *P* value for each fitted data type, i.e. gene expression variation for RNA-seq orLAD size variation for LMNB1 ChIP-seq data. These multiple *P* values are then combined into a one-test Chi-square statistics assuming a Chi-squared distribution with 2 k degrees of freedom (where k is the number of *P* values; here k = 3 for RNA-seq data and k = 2 for LAD data), when all null-hypotheses are true and each *P* value is independent. The combined *P* value is determined by the Chi-square *P* value and was used to determine the significance of oscillating patterns.

### Data access

Lamin B1 ChIP-seq and RNA-seq data have been deposited to the NCBI GEO database and are available under GEO accession No. GSExxxx.

## Abbreviations

ChIP-seq: chromatin immunoprecipitation sequencing
LAD: lamina-associated domain
RNA-seq: RNA-sequencing
TF: transcription factor

## Acknowledgements

We thank Dr. Nolwenn Briand for fruitful discussions. This work was funded by EU Marie Curie Scientia Fellowship FP7-PEOPLE-2013-COFUND No. 609020 (A.B.), the Research Council of Norway (P.C.) and South-East Health Norway (P.C.).

## Author contributions

F.F., A.B. and P.C. designed the study. F.F. did wet-lab experiments. A.B. did bioinformatics analyses. A.B. and P.C. wrote the paper. P.C. supervised the work.

